# How predictable is an individual’s DNA methylation?

**DOI:** 10.64898/2026.11.21.768291

**Authors:** Roei Zucker, Michal Linial, Tom Hope

## Abstract

DNA methylation is largely shared among individuals of the same cell type, yet at specific genomic sites it differs substantially between individuals. Methods that predict methylation are usually evaluated by their accuracy over all sites. Because methylation at most sites is shared, this accuracy cannot show whether a method predicts an individual’s methylation where individuals differ. We present individual-level methylation prediction at sites where individuals differ as a new challenge. Given the reference genome sequence, the methylation of other individuals of the same cell type, and measurements at some sites of a target individual, but not the individual’s genotype, the task is to predict the target individual’s methylation class (unmethylated, intermediate, or methylated) at the remaining sites. We evaluate predictions separately at sites with low and high variation across individuals. We compare four approaches: the average of other individuals, a sequence model (the DNA language model Nucleotide Transformer) fine-tuned on other individuals’ methylation, the same model further fine-tuned on the target individual’s measurements, and a local class-average approach using the target individual’s nearby training-site class labels. Across 32 individuals from six human cell types, the average of other individuals matched or slightly exceeded both sequence models at sites with low variation among individuals. At the most variable sites, the average of other individuals performed worse than a random predictor that assigns classes at the observed class frequencies, whereas both sequence models performed substantially better than the average. Both sequence models also achieved higher mean weighted F1 than the local class-average approach. Fine-tuning on the target individual’s measurements improved predictions more inside regions where other individuals show bimodal methylation (a mix of methylated and unmethylated DNA molecules within one sample) than outside them. Among the most variable sites, it also improved predictions at CpG islands for 25 of the 32 individuals. We conclude that individual-level methylation prediction is a distinct problem that is particularly challenging at sites where individuals differ, and that the benefit of an individual’s own measurements is concentrated in genomic contexts identifiable from other individuals’ methylation and genomic annotations.

## Introduction

DNA methylation is an epigenetic modification that contributes to gene regulation and to the identity of cell types (Kim and Costello, 2017). Methylation patterns differ between cell types and are largely shared among individuals of the same cell type (Loyfer et al., 2023). Yet at specific genomic sites, methylation differs substantially between individuals. Part of this variation is genetic: sequence variants can change local methylation and transcription-factor binding (Banovich et al., 2014; Heyn et al., 2013; Husquin et al., 2018). Environmental and lifestyle exposures also change methylation, sometimes at groups of CpG sites that change together (Lam et al., 2012; Garg et al., 2018). Differences between individuals further depend on cell composition, genomic context, and age (Jacoby et al., 2012; Lokk et al., 2014; Hannum et al., 2013; Talens et al., 2010). These differences are not spread evenly across the genome: some regions are stable across individuals, whereas others differ between individuals in a reproducible way (Bock et al., 2008; Taudt et al., 2016; Gunasekara et al., 2019; Miller et al., 2023).

Computational methods predict DNA methylation from different sources of information. Sequence-based predictors, including CpGenie, INTERACT, and Melody, learn how methylation depends on DNA sequence (Zeng and Gifford, 2017; Zhou et al., 2022; Jin et al., 2026). Genomic language models such as Nucleotide Transformer learn representations of DNA sequence that can be fine-tuned for such tasks (Dalla-Torre et al., 2025). Other methods use measured methylation. DeepCpG predicts missing methylation states in single cells from DNA sequence and from neighboring measured states (Angermueller et al., 2017), and MethylGPT learns representations of methylation profiles (Ying et al., 2024). Imputation methods for bulk data predict methylation at an individual’s unmeasured CpG sites from measured neighboring sites of the same individual, together with genomic features or with other samples measured at the same site (Zhang et al., 2015; Zou et al., 2018). Predicting how individuals differ has proven hard for sequence models: models trained on the reference genome explain little of the variation in gene expression between individuals (Huang et al., 2023; Sasse et al., 2023). SAGE-net trains sequence models on personal genomes and predicts differences in DNA methylation and gene expression between individuals from their genotypes; for methylation, it was trained on array measurements of brain tissue from 634 individuals (Spiro et al., 2026).

Imputation methods use an individual’s own measurements but mainly report accuracy over all sites, and sequence models of personal genomes require the individual’s genotype. These methods therefore leave open how well the methylation of a specific individual can be predicted where individuals differ, without the individual’s genotype, and which information such prediction requires. We pose this question as a new challenge, which we call individual-level methylation prediction (Fig. 1). We call the individual whose methylation we predict the target donor, and the other individuals of the same cell type the reference donors. The input consists of the reference genome sequence, the methylation of the reference donors, and the target donor’s measured methylation at a subset of sites (six-base segments of the genome that contain CpGs), which we call training sites. The output is the target donor’s methylation class (unmethylated, intermediate, or methylated) at the remaining sites. Because methylation at most sites is shared among individuals (Loyfer et al., 2023), accuracy over all sites mainly reflects these shared sites. We therefore evaluate predictions separately at sites with low and high variation among the reference donors.

**Figure 1.**
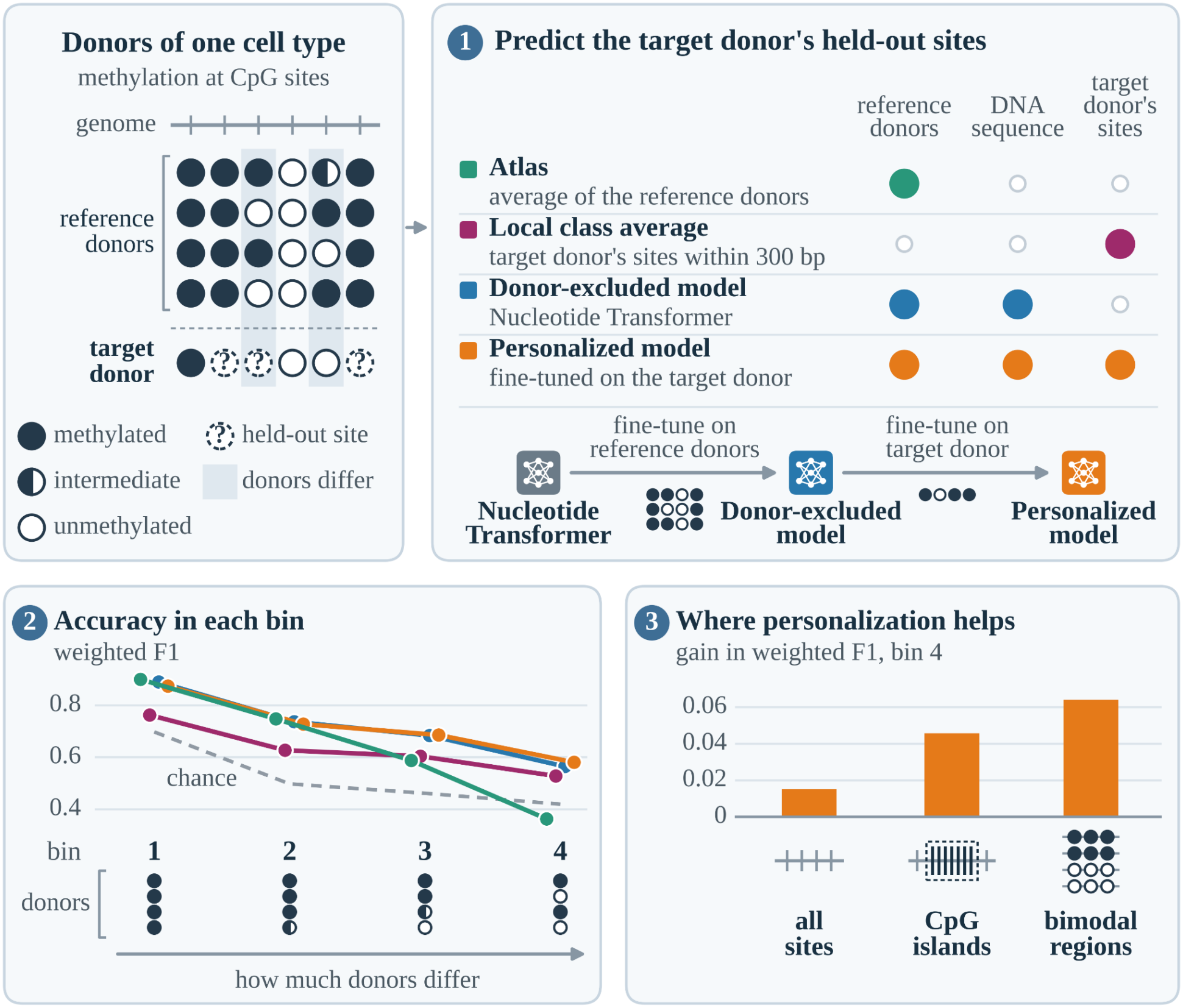
Individual-level methylation prediction. Top left: the task is to predict a target donor’s methylation class at held-out sites, using other donors of the same cell type (reference donors), the DNA sequence, and the target donor’s remaining sites. (1) The four approaches and the information each uses. Below them, Nucleotide Transformer is fine-tuned first on the reference donors, giving the donor-excluded model, and then on the target donor, giving the personalized model. (2) Weighted F1 on the test set, averaged over 32 donors, in four bins that group sites by how much the reference donors differ, from bin 1 (least) to bin 4 (most). The dashed line is a random predictor that follows the target donor’s class frequencies. (3) Personalization gain in bin 4: the personalized model’s weighted F1 minus the donor-excluded model’s.

We compare four approaches that use different combinations of reference-donor measurements, DNA sequence, and target-donor training labels. The first approach, which we call the atlas, predicts the class of the reference donors’ average methylation at each site; like a methylation atlas, it describes the typical methylation of the cell type. The second approach, the donor-excluded model, is Nucleotide Transformer fine-tuned to predict methylation classes from DNA sequence, using only the reference donors’ data. The third approach, the personalized model, is the donor-excluded model further fine-tuned on the target donor’s training sites; we call this second fine-tuning step personalization. The fourth approach, the local class-average approach, predicts each held-out site’s class from nearby training-site class labels of the target donor. We refer to the donor-excluded and personalized models together as the sequence models. We evaluate the approaches on whole-genome bisulfite sequencing data from 32 donors of six human cell types (Loyfer et al., 2023). To find where the target donor’s own measurements help, we compare the approaches within genomic annotations, such as CpG islands, promoters, and candidate cis-regulatory elements (Loyfer et al., 2023; ENCODE Project Consortium, 2020). We also compare them inside and outside bimodal regions, which contain both methylated and unmethylated DNA molecules within one sample (Rosenski et al., 2025). This within-sample bimodality is distinct from differences in methylation level between individuals.

Our results show that individual-level prediction differs from predicting the typical methylation of a cell type. First, at sites with low variation among donors, the atlas matched or slightly exceeded both sequence models. At the most variable sites, it performed worse than a random predictor that assigns classes at the observed class frequencies, while both sequence models performed substantially better than the atlas. Second, personalization gave a small average improvement at the most variable sites, and its benefit differed across cell types and donors. Third, personalization helped more inside regions that are bimodal in reference donors than outside them, with larger mean gains when more reference donors had a bimodal region at the site; among the most variable sites, it also improved predictions at CpG islands.

## Methods

### Individual-level methylation prediction

We define the task for one target donor at a time. The input has three parts: the DNA sequence of the human reference genome (hg38); the methylation of the reference donors, which are the other donors of the same cell type; and the target donor’s measured methylation at a subset of sites, which we call training sites. The output is the target donor’s methylation at the remaining measured sites, which we call held-out sites. The main evaluation tests prediction of held-out target-donor labels within genomic regions used to train the donor-excluded model on reference-donor labels. We divided the held-out sites further into a validation set, which we used for development and to monitor training, and a test set, on which we report the main results. Nucleotide Transformer reads DNA in consecutive six-base units (tokens). We therefore define a site as a token that contains at least one measured CpG; a site can contain more than one CpG. For each site, we predict one of three methylation classes: unmethylated (methylation level of at most 0.2), intermediate (above 0.2 and below 0.8), or methylated (at least 0.8).

Because methylation at most sites is shared among donors, performance over all sites mainly reflects these sites. We therefore report performance separately in four variability bins, which group sites by the standard deviation of their methylation among the reference donors, from bin 1 (lowest) to bin 4 (highest). The bins use only the reference donors, so the target donor’s measurements do not affect bin assignment.

### Data

We used whole-genome bisulfite sequencing data of human cell types from Loyfer et al. (2023). We selected the cell types with at least four donors. White-blood-cell samples (n = 23) were excluded to limit potential effects of cell composition on methylation profiles (Reinius et al., 2012). The 36 purified blood-cell samples belonged to cell-type subgroups with fewer than four donors each and therefore did not meet the inclusion criterion. The resulting data contain 32 donors of six cell types: bladder epithelial cells (5 donors), breast basal epithelial cells (4), cortex neurons (8), cardiomyocytes (5), heart fibroblasts (4), and liver hepatocytes (6). We used the processed methylation tracks released by Loyfer et al., in which the methylation level of a CpG is the fraction of sequenced DNA fragments in which it is methylated. We divided the genome into non-overlapping 5,400-bp windows; each window, 900 six-base tokens long, is one model input. In preliminary experiments, 5,400-bp windows gave better predictions than shorter windows. The methylation level of a site is the mean methylation level of the measured CpG positions in it. Appendix A gives the remaining preprocessing details.

### Prediction approaches

We compared four prediction approaches: the atlas, donor-excluded model, personalized model, and local class-average approach. A class-frequency random predictor provided a chance-level reference.

#### Atlas

The atlas predicts the class of the reference donors’ average methylation at each site. We averaged the reference donors’ methylation levels at each CpG measured in all of them, averaged these values over the CpG positions of each site, and assigned a class with the thresholds above. Sites not measured in all reference donors have no atlas prediction; comparisons that include the atlas use only sites predicted by all approaches.

#### Donor-excluded model

This model predicts methylation classes from DNA sequence after training on the reference donors only. We started from the pretrained Nucleotide Transformer with 2.5 billion parameters, trained on the genomes of many species (Hugging Face model InstaDeepAI/nucleotide-transformer-2.5b-multi-species; Dalla-Torre et al., 2025). We added a classification head that scores the three classes for each token, and the model predicts the class with the highest score. For each target donor, we fine-tuned this model on the classes of all measured sites of the reference donors in windows on chromosomes 1–5 and 18–22, with training configured for 12 epochs, and used the checkpoint after epoch 10. We selected epoch 10 using performance on the validation set.

#### Personalized model

This model adds the target donor’s own measurements. We initialized it from the target donor’s donor-excluded model at epoch 10 and fine-tuned all of its parameters on the target donor’s training sites, with a higher learning rate and with class rebalancing (Appendix A). Fine-tuning was restricted to chromosomes 1–5 and 18–22. We trained each personalized model for three epochs, the training duration available for these experiments, and used the final checkpoint.

#### Class-frequency random predictor

This predictor assigns each site a class at random, with the frequencies of the target donor’s true classes in the evaluated set of sites. Because it uses the true class frequencies, it indicates chance-level performance rather than a usable method. We computed its weighted F1, defined below, from the expected confusion counts.

#### Local class-average approach

This approach used only the target donor’s training-site class labels. For each evaluated site, we calculated the unweighted mean of neighboring training labels, encoded as 0, 1 and 2, within a fixed genomic radius on the same chromosome. Means below 0.5 were assigned to the unmethylated class, means from 0.5 to below 1.5 to the intermediate class, and means of at least 1.5 to the methylated class. When no training labels occurred within the radius, we used the nearest training site’s class; chromosomes without training labels used the donor’s most frequent training class. A radius of 300 bp in either direction was selected by maximizing equal-donor mean weighted F1 on the validation set and then fixed for all donors in testing.

### Training, validation, and test sites

We divided each target donor’s measured sites in two steps. The target donor’s training, validation and test windows came from chromosomes 1–5 and 18–22. The same DNA windows could also occur in donor-excluded training, paired with the reference donors’ methylation classes. First, within each 5,400-bp window, we assigned about 80% of the measured sites to training and held out the remaining 20%. The training version and the held-out version of a window both contain the window’s full DNA sequence; they differ only in which sites’ classes they include. Second, we assigned the held-out version of each window either to the validation set (about 20% of windows) or to the test set (about 80% of windows). A test window can therefore also appear in the personalized model’s training, with the target donor’s classes at different sites.

### Variability bins

For each target donor, we computed the sample standard deviation of methylation among the reference donors at each CpG measured in all of them, and averaged these values over the CpG positions of each site. We assigned sites to four bins using cutoffs at one, two, and three fifths of the largest standard deviation of any CpG: bin 1 contains the least variable sites and bin 4 the most variable. Bin 4 combines the two highest fifths, because the highest fifth alone contains few sites. Sites without measurements in all reference donors have no bin. Appendix A gives details.

### Performance measures

We measured performance with weighted F1: the F1 score of each class, averaged over the three classes with weights equal to each class’s share of the evaluated sites. We computed weighted F1 separately for each target donor and each evaluated set of sites, such as the bin-4 test sites of one donor, and then averaged over donors with equal weights. The personalization gain of a donor is the personalized model’s weighted F1 minus the donor-excluded model’s weighted F1, computed on the same sites.

### Genomic annotations

We evaluated predictions within CpG contexts (islands, shores, shelves and open sea), promoters, gene bodies, and ENCODE candidate cis-regulatory elements (cCREs), including a regulatory union. Sites were assigned by overlap; each donor–annotation–bin combination required at least 50 test sites. CpG contexts were mutually exclusive, while other annotations could overlap. These analyses used existing predictions without retraining. Appendix A gives annotation sources, versions and interval definitions.

### Bimodal regions and cross-donor support

Rosenski et al. (2025) annotated bimodal regions, which contain both methylated and unmethylated DNA molecules within one sample, separately for each sample of the Loyfer et al. (2023) data. A site is inside a bimodal region if it overlaps a region annotated as bimodal in at least one reference donor. Support from the target donor was excluded, but region boundaries were retained from the published multi-sample catalog, which included target-donor samples where available. The boundaries were not rebuilt separately for each evaluated donor. The cross-donor support of a site is the number of reference donors with a bimodal region that overlaps the site, grouped as zero, one, or at least two. Sites with zero reference-donor support were outside bimodal regions. Appendix A describes coordinate mapping, unmapped sites and the sensitivity analysis.

### Chromosomes 6–17

We also evaluated the two sequence models on chromosomes 6–17, which were excluded from supervised methylation training at both stages. Appendix A describes chromosome- and cell-type-exclusion checks.

#### Donor-matching control

On chromosomes 6–17, we compared each donor’s matched model with models for other donors of the same cell type, using identical evaluation sites, separately in bins 3 and 4. We performed this comparison for both personalized and donor-excluded models; Appendix A gives the scoring and aggregation procedures.

### Uncertainty and multiple testing

We computed 95% confidence intervals using a percentile bootstrap over donors (20,000 resamples), pairing scores within donors for comparisons and resampling within cell types for cell-type summaries. Exact two-sided paired donor sign tests assessed consistency of differences across donors, excluding ties. We applied Benjamini–Hochberg (BH) correction within the specified comparison families; the regulatory screen comprised 22 comparisons across bins 3 and 4. Confidence intervals were not adjusted for multiple comparisons. Appendix A gives the testing families and implementation details.

### Data and code availability

Processed source methylation tracks are available through GEO accession GSE186458 (Loyfer et al., 2023). Code is available at https://github.com/RoeiZucker/individual-methylation-prediction.

## Results

### Sequence models outperform the atlas at variable sites

For the four approaches and the random predictor alike, performance decreased as the variability among the reference donors increased (Fig. 2). Bins 1 and 2 contained 95.61% of the test donor–site observations assigned to a variability bin, whereas bins 3 and 4 contained 3.79% and 0.60%, respectively. In bins 1 and 2, which contain the least variable sites, the atlas had slightly higher mean weighted F1 than both sequence models. In bins 3 and 4, both sequence models outperformed the atlas, and in bin 4 the atlas performed worse than the class-frequency random predictor. Bin 4 contains, by construction, the sites where the reference donors differ most. At such a site, the average of the reference donors can fall in the intermediate class even when every donor is clearly methylated or unmethylated, so the average matches none of them. The sequence models, in contrast, are trained on each reference donor’s classes rather than on their average, so their predictions are not tied to the class of the average; like the atlas, the donor-excluded model uses no data from the target donor in training. On average, personalization increased weighted F1 over the donor-excluded model by 0.0157 in bin 4 (95% donor-bootstrap CI, 0.0029–0.0286), changed it little in bin 3, and slightly reduced it in bins 1 and 2. The local class-average approach achieved mean weighted F1 values of 0.762, 0.626, 0.603, and 0.527 in bins 1–4, respectively.

**Figure 2.**
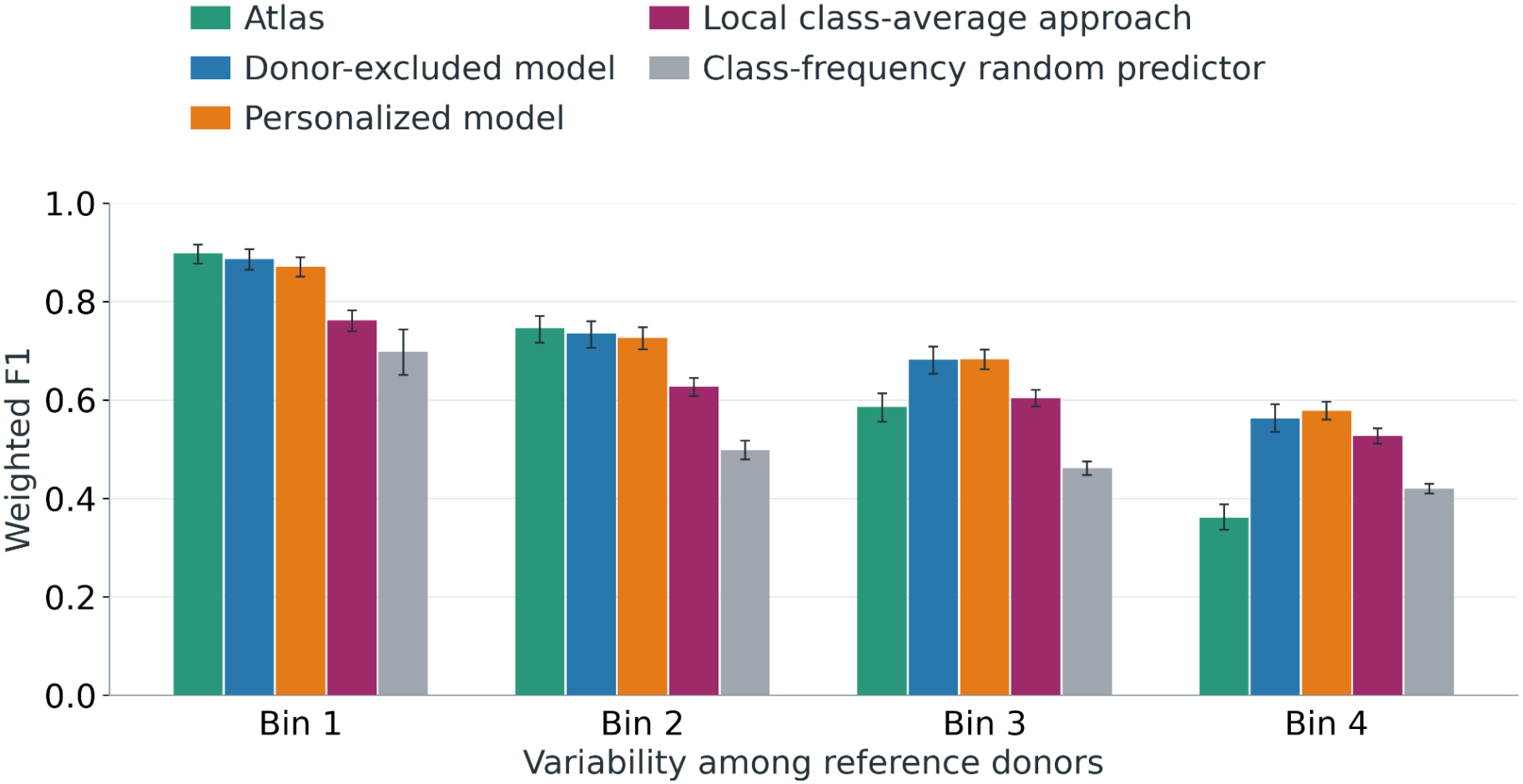
Prediction performance across methylation-variability bins. Bars show weighted F1 averaged equally over the 32 donors for the atlas, donor-excluded model, personalized model, local class-average approach, and class-frequency random predictor. Whiskers show 95% bootstrap confidence intervals over donors.

### Personalization gains vary across cell types and donors

The personalization gain differed across cell types and donors (Fig. 3). In bins 3 and 4, both sequence models outperformed the atlas and the class-frequency random predictor in every cell type. The mean personalization gain was positive in both bins for bladder epithelial cells, breast basal epithelial cells, and liver hepatocytes, and it was larger in bin 4 for the latter two. Heart fibroblasts had small positive mean gains in both bins. Cortex neurons had negative mean gains in both bins, and cardiomyocytes had a negative mean gain in bin 3 and a slightly positive one in bin 4. In bin 4, the mean gain ranged from ™0.0099 in cortex neurons to +0.0464 in liver hepatocytes. With 4 to 8 donors per cell type, these estimates are imprecise. Across all cell types, personalization improved weighted F1 for 15 of the 32 donors in bin 3 and 21 of the 32 in bin 4; in neither bin did a sign test distinguish this from an even split. Personalization also shifted performance between classes: in bins 3 and 4, it increased the mean F1 of the unmethylated and intermediate classes and reduced the mean F1 of the methylated class (Figure A1).

**Figure 3.**
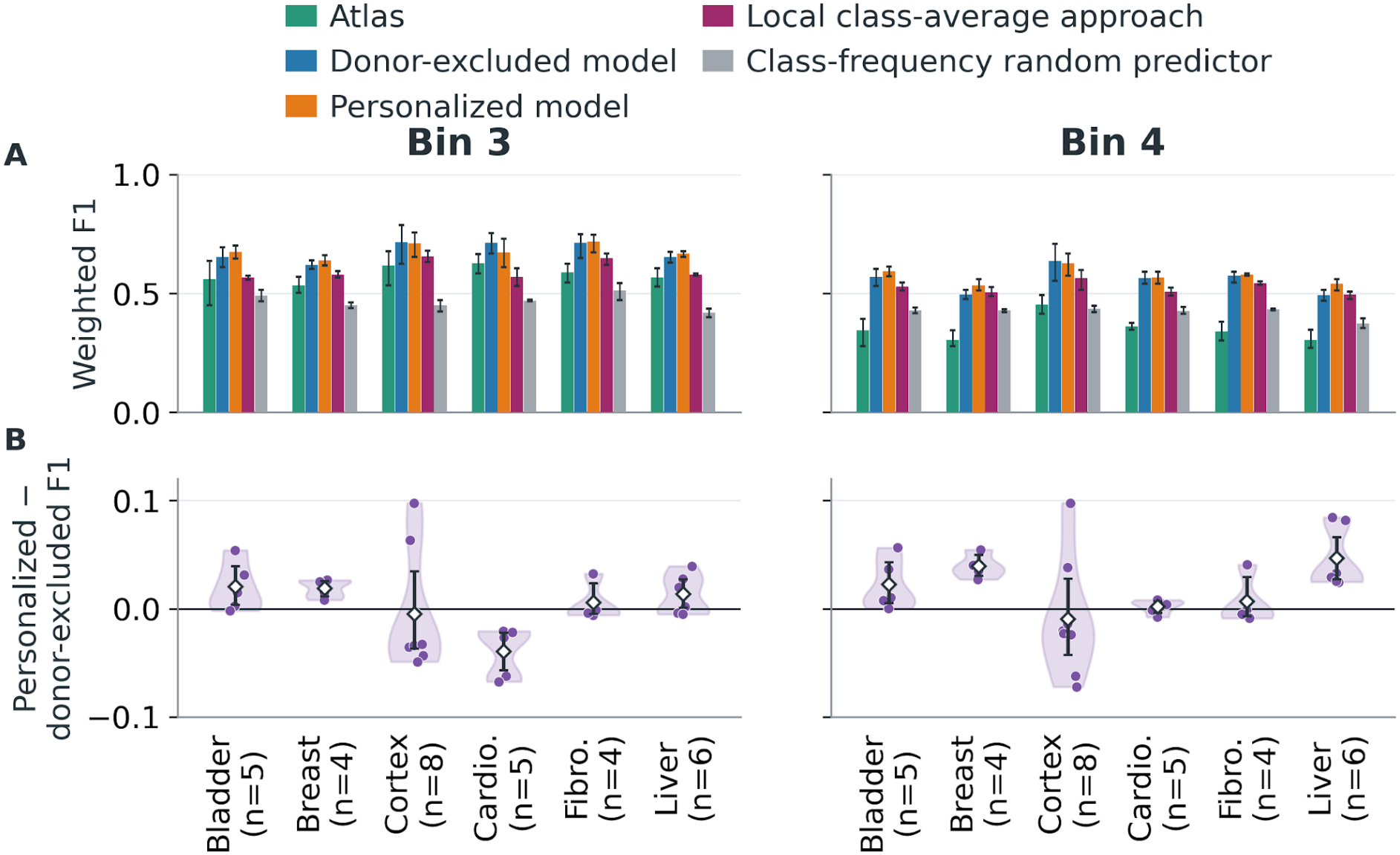
Performance and personalization gains across cell types in bins 3 and 4. (A) Weighted F1 averaged equally over donors for the atlas, donor-excluded model, personalized model, local class-average approach, and class-frequency random predictor. (B) Personalization gains (personalized minus donor-excluded weighted F1); dots represent donors and violins show their distributions. Diamonds mark mean gains; whiskers show 95% bootstrap confidence intervals over donors. Results include all 32 donors, with 4–8 per cell type. Cardio. and Fibro. denote cardiomyocytes and heart fibroblasts.

### Personalization gains on chromosomes 6–17

On chromosomes 6–17, excluded from supervised methylation training at both stages, the mean personalization gain was positive in bins 3 and 4 across all 32 donors and remained positive in the chromosome- and cell-type-exclusion checks (Appendix A; Table A3).

In an exploratory donor-matching control on these chromosomes, matched personalized models achieved higher mean weighted F1 than models personalized for other donors of the same cell type in both bins. In bin 4, the matching advantage was positive for 24 of 32 donors (BH-adjusted paired donor sign test, q = 0.028). The corresponding donor-excluded models had mean matching gaps of ™0.0018 in bin 3 and ™0.0015 in bin 4, with 95% confidence intervals including zero in both bins (Appendix A).

### Personalization improves predictions at CpG islands among the most variable sites

We compared the approaches within genomic annotations in bins 3 and 4 (Fig. 4). At CpG islands in bin 4, personalization raised the mean weighted F1 by 0.0457 and improved predictions for 25 of the 32 donors (two-sided sign test, p = 0.0021; Benjamini–Hochberg q = 0.046). This was the only one of the 22 comparisons that passed the Benjamini–Hochberg correction. CpG islands and promoter-like cCREs in bin 3, and CpG shores in bin 4, showed gains with p < 0.05 that did not pass the correction. In most annotations, personalization improved predictions for more donors than it worsened.

**Figure 4.**
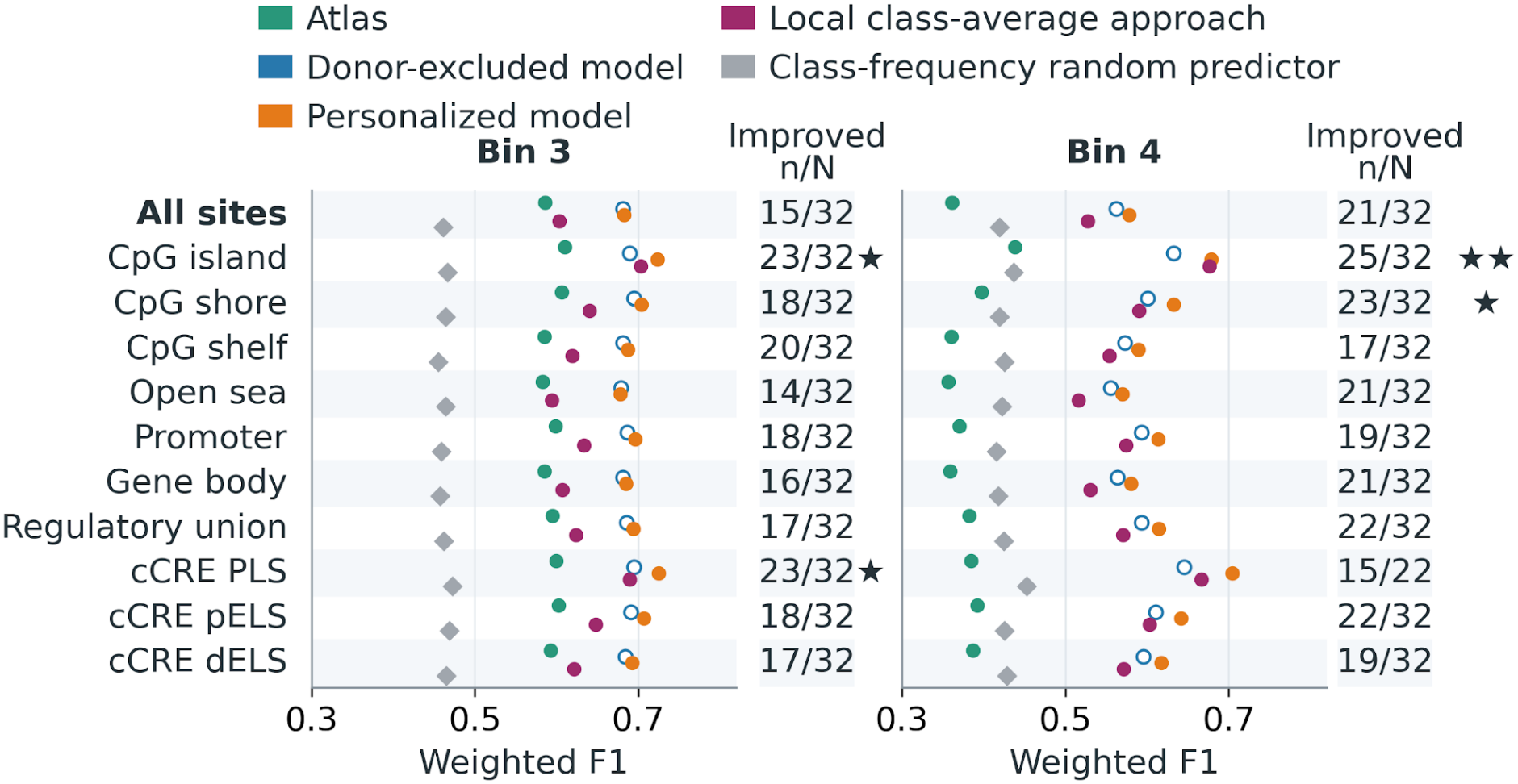
Exploratory regulatory screen in bins 3 and 4. Points show weighted F1 averaged equally over eligible donors. The local class-average approach is shown in magenta. Counts give the number of donors whose personalized model outperformed their donor-excluded model, out of all eligible donors. Stars require a positive mean personalization gain: one star marks a donor sign-test p < 0.05 that does not pass Benjamini–Hochberg correction; two stars mark q < 0.05 after correction across 22 comparisons. Each donor was eligible within an annotation and bin if at least 50 test sites were available. CpG contexts are mutually exclusive; other annotations can overlap.

### Personalization gains are larger in bimodal regions

Personalization gains were larger inside bimodal regions than outside them (Fig. 5; Table A2). A site counted as inside if it overlapped a region that was bimodal in at least one reference donor; support from the target donor was excluded. Inside bimodal regions, the personalized model had higher mean weighted F1 than the atlas and donor-excluded model in both bins 3 and 4, and personalization improved predictions for 25 of 32 donors in bin 3 and 26 of 32 in bin 4. Outside bimodal regions, the mean gains were smaller, and their 95% confidence intervals included zero in both bins. Within donors, the gain inside bimodal regions exceeded the gain outside them for 31 of 32 donors in bin 3 and 28 of 32 in bin 4 (exact two-sided paired donor sign tests, Benjamini–Hochberg q < 0.001 in both bins; Table A2). We compared personalized predictions with a local class-average approach using nearby target-donor training-site class labels. The personalized model achieved higher mean weighted F1 across bins 3 and 4 overall. Within bimodal regions, mean weighted F1 was 0.726 versus 0.703 in bin 3, with a positive paired difference supported by both the donor-bootstrap confidence interval and the BH-adjusted donor sign test (q < 0.001). In bin 4, mean weighted F1 was also higher for the personalized model (0.677 versus 0.660), but the paired difference confidence interval included zero, so an advantage was not established.

**Figure 5.**
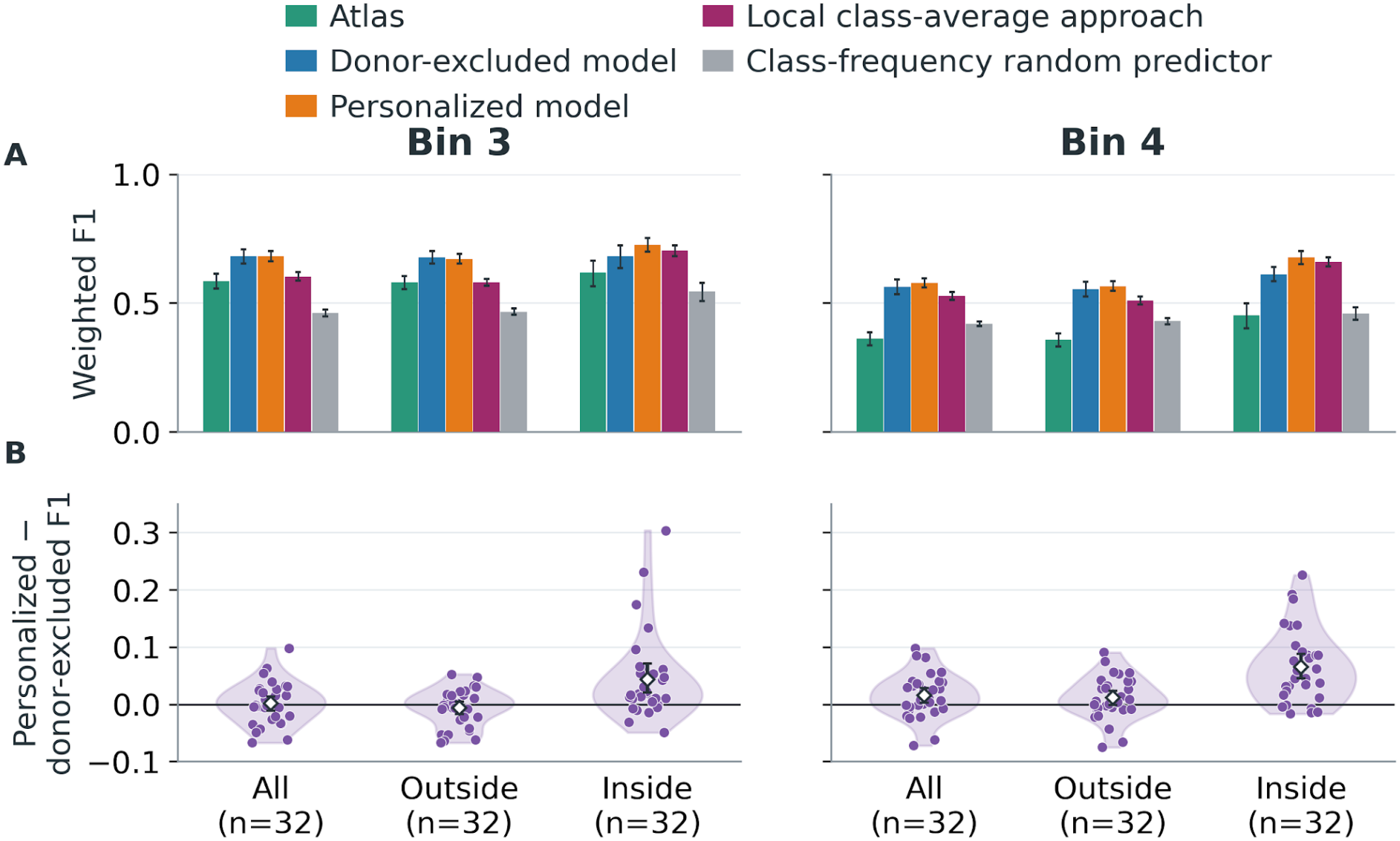
Performance and personalization gains inside and outside annotated bimodal regions in bins 3 and 4. (A) Equal-donor mean weighted F1 for the four approaches and class-frequency random predictor. (B) Personalized-minus-donor-excluded F1; dots represent donors, violins their distributions, and diamonds the means. Whiskers show 95% donor-bootstrap confidence intervals. All 32 donors contribute to each group. Inside denotes support from at least one reference donor; outside denotes none. “All” combines both groups.

### Mean gains are larger when more reference donors have a bimodal region at the site

We then grouped sites by cross-donor support, the number of reference donors with a bimodal region that overlaps the site: zero (outside bimodal regions), one, or at least two (Table 1). The mean personalization gain was larger with more supporting donors in both bins. In bin 3, the gain was ™0.007 with no supporting donor, +0.021 with one, and +0.057 with at least two. In bin 4, the corresponding gains were +0.011, +0.053, and +0.080. Paired donor comparisons supported larger gains in each higher-support group in bin 3. In bin 4, both supported groups had larger gains than the zero-support group, whereas the difference between one and at least two supporting donors was not established (Table 1).

**Table 1.** Paired comparisons of personalization gains between cross-donor-support groups.

| Bin | Support groups compared | Mean gain difference [95% CI] | p-value | BH q-value |
|---|---|---|---|---|
| 3 | 1 versus 0 | +0.028 [+0.016, +0.041] | 1.93 × 10<sup>−5</sup> | 3.86 × 10<sup>−5</sup> |
| 3 | ≥2 versus 1 | +0.036 [+0.019, +0.054] | 1.93 × 10<sup>−5</sup> | 3.86 × 10<sup>−5</sup> |
| 3 | ≥2 versus 0 | +0.064 [+0.043, +0.088] | 4.66 × 10<sup>−10</sup> | 2.79 × 10<sup>−9</sup> |
| 4 | 1 versus 0 | +0.042 [+0.022, +0.065] | 0.00210 | 0.00252 |
| 4 | ≥2 versus 1 | +0.027 [−0.001, +0.054] | 0.110 | 0.110 |
| 4 | ≥2 versus 0 | +0.069 [+0.047, +0.093] | 0.000113 | 0.000170 |

Differences are gains in the higher-support group minus those in the lower-support group, paired by donor. p-values are exact two-sided donor sign tests, excluding ties; q-values apply BH correction across all six comparisons. CIs are unadjusted 95% paired donor-bootstrap intervals (20,000 resamples; seed 20261104).

## Discussion

We presented individual-level methylation prediction as a new challenge: predicting the methylation of a specific individual from the reference genome sequence, from the methylation of other individuals of the same cell type, and from measurements at some of the individual’s own sites. Evaluating predictions separately by how much sites vary among donors showed where the challenge lies. Where donors agree, neither sequence model improved on the average of the reference donors. Where donors disagree, sequence models trained on the reference donors performed substantially better than this average, which fell below the class-frequency random predictor at the most variable sites. Fine-tuning on the target donor’s own measurements added gains inside regions that are bimodal in reference donors and at CpG islands among the most variable sites.

Accuracy over all sites hides these differences, because sites where donors agree dominate it.

Personalization helped more inside regions that are bimodal in reference donors than outside them, and mean gains were larger when more reference donors had a bimodal region at the site. Rosenski et al. (2025) identified such regions in their study of imprinted and allele-specific methylation. In regions with sequence-dependent allele-specific methylation, a donor’s methylation depends on which alleles the donor carries. Our models read only the reference genome, not the target donor’s genotype, so for such regions the donor’s own measurements are the models’ only source of information about the donor’s alleles. This may explain why these measurements help more inside bimodal regions than outside them. Bimodal regions contained more intermediate and fewer methylated target labels than outside regions, so differences in class composition may contribute to the larger personalization gains. Regions with different cross-donor support may also differ in other properties, so our results do not establish why the gains differ. They do show that the average of other individuals misses patterns in how methylation differs between individuals, and that a model can learn part of an individual’s methylation from partial measurements of that individual.

The four approaches use different combinations of reference-donor measurements, DNA sequence, and target-donor training labels. Comparing the donor-excluded model with the atlas measures what a sequence model trained on the reference donors adds to their average; comparing the personalized model with the donor-excluded model measures what fine-tuning on the target donor’s own measurements adds. Personalization gains differed across cell types and donors, so an average over all donors can hide both gains and losses. Our local class-average comparison assessed prediction from nearby target-donor class labels, using neighboring methylation information as imputation methods do (Zhang et al., 2015; Zou et al., 2018). Within bimodal regions, the personalized model showed a statistically supported advantage in bin 3. In bin 4, it achieved higher mean weighted F1, although the paired confidence interval included zero and the donor sign test was not significant. The CpG-island finding demonstrates improvement over the donor-excluded model, but does not establish an advantage over the local class-average approach in bin 4. Improvements over the donor-excluded model therefore do not imply a consistent advantage over the local class-average approach. The same task and evaluation also accommodate further approaches. One would use the most common class among the reference donors, or the reference donor most similar to the target donor, instead of their average. Another would add the target donor’s genotype, as models trained on personal genomes do (Spiro et al., 2026). Comparing these approaches at sites where donors differ would show which information individual-level prediction needs most.

### Limitations

Our data contain 32 donors of six cell types, with 4 to 8 donors per cell type, which limits the precision of the cell-type estimates and the generality of the results. Mean personalization gains were positive in both bins for bladder epithelial cells, breast basal epithelial cells, and liver hepatocytes, small for heart fibroblasts, and negative in at least one bin for cardiomyocytes and cortex neurons. Larger cohorts are needed to tell whether these differences reflect the cell types or the donors sampled. We excluded blood cells and all cell types with fewer than four donors. The personalization gain combines the effect of the target donor’s data with the effect of further training, which used a higher learning rate and class rebalancing. The donor-matching control showed positive mean matching advantages on chromosomes 6–17. However, models personalized for other donors started from donor-excluded models that had already been trained on the target donor’s labels from chromosomes 1–5 and 18–22. This difference in training membership limits how cleanly the control isolates the effect of donor matching. We trained the personalized models for only three epochs. Finally, our models did not use the donors’ genotypes, which affect methylation at many sites.

Evaluation at variable sites reveals differences hidden by overall accuracy. Sequence models outperform the atlas at these sites, while personalization adds gains in reference-donor bimodal regions and CpG islands among the most variable sites. Larger cohorts, more cell types, and integration of donor genotypes are natural next steps.

## Acknowledgments

This research was made possible through a GPU compute resource grant funded by the Association of University Heads, the Council for Higher Education and the AI Research Compute Center.

