## Supplemental Data 1 for "How predictable is an individual’s DNA methylation?"

### Appendix A. Implementation details

**Preprocessing.** We obtained DNA sequences from the hg38 reference genome and methylation levels from the donor-specific processed BigWig tracks released by Loyfer et al. (2023). According to their processing records, reads were aligned with bwa-meth, duplicates were marked with Sambamba, a mapping-quality threshold of 10 was applied, and BigWig tracks were generated with default beta2bw parameters. We applied no additional read-depth threshold. We converted BigWig values to float16 before expanding them to single bases. Before averaging within a token, we masked a measured first base when the token's second base was unmeasured. Each token's methylation level is the mean of its measured per-base values, stored as float32. Tokens without measurements and the model's special token were excluded from the training loss and from evaluation. We discarded windows with ambiguous bases or with incorrect sequence or label lengths.

**Class thresholds.** Both target labels and atlas predictions were classified as unmethylated (≤0.2), intermediate (>0.2 and <0.8), or methylated (≥0.8).

**Training data filters.** The donor-excluded models used all measured sites of the reference donors in each window, with no sites held out. For the personalized models, we randomly masked excess training sites so that no class had more than four times as many sites as the smallest class, and we removed windows left without training sites. Both training stages required at least ten training sites per window. We applied neither the class balancing nor the ten-site filter to the validation and test sets, which kept every window with at least one held-out site.

**Training.** Both stages updated all parameters of the Transformer and of the classification head. We used the PyTorch implementation of AdamW ($\beta1 = 0.9$, $\beta2 = 0.999$, $\varepsilon = 10^{-8}$), a linear learning-rate schedule without warmup, weight decay 0.15, gradient clipping at a maximum norm of 1, bfloat16 precision, and random seed 42. The batch size was 14 on one GPU, without gradient accumulation. The learning rate was $1\times10^{-6}$ for the donor-excluded models and $3\times10^{-5}$ for the personalized models. We used Hugging Face Transformers 4.41.0. For the personalized models, we used the checkpoint after epoch 3 without selecting a best checkpoint.

**Epoch selection for the donor-excluded models.** Epoch 10 was selected using validation-set performance and used consistently across all donor-excluded models.

**Variability bins.** We joined the reference donors' measurements on identical CpG coordinates and computed the sample standard deviation at each CpG measured in all reference donors. The variability of a site is the mean standard deviation of the CpGs whose annotated start lies between one base before and five bases after the start of the site's token (offsets −1 to +5); the atlas uses the same assignment of CpGs to sites. Let M be the largest CpG-level standard deviation among the target donor's reference donors, before averaging within sites. Bins 1–4 are [0, M/5], (M/5, 2M/5], (2M/5, 3M/5], and (3M/5, M]. Bin 4 combines the two highest fifths: in the test set, the highest fifth contained a median of 728 sites per donor, compared with 8,543 in the fourth fifth. We computed the bin-4 F1 of each donor from the combined sites.

**Table A1. Distribution of binned test donor–site observations.**

| Bin | Donor–site observations | Share of binned test observations |
|---|---:|---:|
| 1 | 45,442,697 | 79.85% |
| 2 | 8,967,720 | 15.76% |
| 3 | 2,157,751 | 3.79% |
| 4 | 340,913 | 0.60% |
| **Total** | **56,909,081** | **100%** |

Counts are pooled donor–site observations across 32 donors, not unique genomic sites. Percentages use the 56,909,081 test observations assigned to a variability bin; observations without a bin assignment are excluded.

**Genomic annotations.** We used UCSC cpgIslandExt (hg38), GENCODE v50 (GRCh38.p14), and the ENCODE SCREEN Registry v4. A site belongs to an annotation when any of its six bases overlaps the annotation. We assigned CpG contexts hierarchically, in the order island, shore (within 2 kb of an island), shelf (2–4 kb from an island), and open sea (all remaining sites), so each site has exactly one context. Promoters span 2 kb upstream to 500 bp downstream of transcription start sites, accounting for the gene's strand; gene bodies span whole genes. We report promoter-like (PLS), proximal enhancer-like (pELS), and distal enhancer-like (dELS) cCREs separately. The regulatory union combines promoters with all SCREEN cCRE classes. We merged overlapping intervals within each annotation before assigning sites. Apart from the CpG contexts, annotations can overlap. In analyses of genomic annotations, we included a donor only if the annotation contained at least 50 of that donor's test sites in the bin.

**Mapping to bimodal regions.** We mapped sites from hg38 to hg19 with the UCSC hg38-to-hg19 liftOver chain. We accepted a mapping only when both ends of the site mapped uniquely to the same chromosome and strand and formed an interval of at most 20 bp. Each reference donor counts once toward the cross-donor support of a site, even when several of its bimodal regions overlap the site. Sites with failed or rejected mappings were counted as outside bimodal regions in the main analysis. Excluding them in a sensitivity analysis removed 23,704 donor–site observations in bin 3 and 3,283 in bin 4. Mean personalization gains outside bimodal regions changed by less than 0.0002 F1 in both bins, and their 95% donor-bootstrap confidence intervals still included zero. Gains inside bimodal regions were unchanged.

**Chromosomes 6–17.** These analyses include all 32 donors. To test whether personalization gains carry over to chromosomes excluded from supervised methylation training at both stages, we also evaluated the two sequence models on chromosomes 6–17. For each donor, we averaged the personalization gains of the individual chromosomes, and then averaged over donors. To test whether single chromosomes or cell types drove the result, we repeated this calculation after omitting each chromosome, and after omitting all donors of each cell type, one at a time. These analyses used existing predictions; we did not retrain any model.

**Chromosome-exclusion results.** On chromosomes 6–17, which were excluded from supervised methylation training at both stages, the mean personalization gain was positive in both bins 3 and 4. In bin 3, this differs from the main results, where personalization changed performance little. This comparison includes all 32 donors. The gains stayed positive, with 95% confidence intervals above zero, when we omitted any one of the 12 chromosomes or all donors of any one cell type (Table A3).

**Donor-matching control.** We assessed donor matching separately in bins 3 and 4 on chromosomes 6–17. For each target donor, we compared the model personalized for that donor with models personalized for other donors of the same cell type, using only evaluation sites available for all compared models and the target donor's true class labels. Confusion counts were pooled across chromosomes before calculating each model's weighted F1. The matching advantage was the matched model's F1 minus the mean F1 of the other donors' models; these advantages were averaged equally across donors. We repeated the comparison using the corresponding donor-excluded models. All comparisons used existing predictions without additional training.

**Donor-matching results.** Matched personalized models achieved higher mean weighted F1 in bins 3 and 4, with mean matching advantages of 0.014 and 0.011, respectively. In bin 4, the advantage was positive in 24 of 32 donors and supported by a BH-adjusted paired donor sign test ($q = 0.028$). The corresponding donor-excluded models had mean matching gaps of −0.0018 (95% donor-bootstrap CI, −0.0041 to 0.0006) in bin 3 and −0.0015 (95% donor-bootstrap CI, −0.0032 to 0.0003) in bin 4.

**Uncertainty and multiple testing.** We computed 95% confidence intervals for mean performance and mean personalization gains with a percentile bootstrap over donors (20,000 resamples). For gains, we resampled donors and kept each resampled donor's scores for both sequence models together, so each gain is computed within a donor; for cell-type summaries, we resampled donors within each cell type. In the annotation analysis, we tested whether personalization improved predictions for more donors than it worsened, using an exact two-sided sign test that excludes ties. We applied Benjamini–Hochberg (BH) correction across the 22 comparisons of the personalized and donor-excluded models: 11 site groups (all sites, the four CpG contexts, promoters, gene bodies, the regulatory union, and the three cCRE classes) in each of bins 3 and 4. The local class-average and donor-matching comparisons also used exact two-sided paired donor sign tests, excluding ties. Donor-matching p-values were BH-adjusted across four comparisons: two model stages in each of bins 3 and 4. For the local class-average approach, the reported q-values retain BH correction from the broader control screen of 190 weighted-F1 comparisons. We did not adjust confidence intervals for multiple comparisons.

For the direct comparison of personalization gains inside versus outside bimodal regions, we applied exact two-sided paired donor sign tests to the within-donor gain differences, excluding ties, with Benjamini–Hochberg correction across the two comparisons in bins 3 and 4.

Exploratory differences in personalization gain between cross-donor-support groups were evaluated using all three paired group contrasts in each of bins 3 and 4. Exact two-sided donor sign tests excluded ties, with BH correction across the six comparisons. Unadjusted 95% confidence intervals for the mean paired gain differences used 20,000 donor-bootstrap resamples with seed 20261104.

**Bootstrap seeds.** The primary performance and personalization-gain summaries used random seed 20261014; the local class-average and donor-matching comparisons used seed 20261102.

##### Table A2. Prediction performance and personalization gains by regional bimodality

###### A. Performance and personalization gains by regional bimodality. Numerical companion to main Figure 5; recurrence remains main Table 1.

| Bin | Sites | E10 F1 | E3 F1 | Gain [95% CI] | Improved | p | q |
|---|---|---|---|---|---|---|---|
| 3 | All | 0.6813 | 0.6826 | +0.0013 [−0.0109, +0.0139] | 15/32 | 0.860 | 0.860 |
| 3 | Outside | 0.6785 | 0.6718 | −0.0067 [−0.0176, +0.0041] | 14/32 | 0.597 | 0.597 |
| 3 | Inside | 0.6828 | 0.7263 | +0.0435 [+0.0203, +0.0709] | 25/32 | 0.00210 | 0.00420 |
| 4 | All | 0.5625 | 0.5782 | +0.0157 [+0.0029, +0.0286] | 21/32 | 0.110 | 0.147 |
| 4 | Outside | 0.5543 | 0.5655 | +0.0112 [−0.0012, +0.0236] | 20/32 | 0.215 | 0.287 |
| 4 | Inside | 0.6119 | 0.6773 | +0.0654 [+0.0450, +0.0879] | 26/32 | 0.000535 | 0.00214 |

Values are equal-donor mean support-weighted F1. Gain is personalized E3 minus donor-excluded E10, calculated within donors on the same prediction rows. Brackets show nominal 95% percentile donor-bootstrap confidence intervals (20,000 resamples). Improved donors have gain > 0. All rows include 32 donors. Inside requires a published bimodal annotation supported by at least one other same-tissue donor, excluding the evaluated donor; outside is its complement. All includes both subsets, so groups are not independent cohorts. P-values are exact two-sided donor sign tests, excluding ties. BH-adjusted q-values retain the existing families: four whole-bin tests for All, and four inside/outside subset tests across bins 3 and 4. Confidence intervals quantify mean gains; sign tests assess the balance of positive and negative donor gains.

###### B. Paired inside–outside gain contrasts

| Bin | Inside−outside gain [95% CI] | Larger inside | p | q |
|---|---|---|---|---|
| 3 | +0.0502 [+0.0314, +0.0731] | 31/32 | $1.54\times10^{-8}$ | $3.07\times10^{-8}$ |
| 4 | +0.0541 [+0.0381, +0.0716] | 28/32 | $1.93\times10^{-5}$ | $1.93\times10^{-5}$ |

The contrast is each donor's inside-region gain minus that donor's outside-region gain, then averaged equally across donors. Brackets show nominal 95% donor-bootstrap confidence intervals (20,000 resamples). Larger-inside counts refer to positive paired contrasts; they are distinct from the counts improving over E10. These intervals are not multiplicity-adjusted. P-values are exact two-sided paired donor sign tests, excluding ties; q-values use BH correction across the two inside–outside contrasts in bins 3 and 4.

**Table A3. Sensitivity of personalization gains to chromosome and cell-type exclusions**

Results use chromosomes 6–17, excluded from supervised methylation training at both stages, in the 32-donor cohort. Gains are personalized-model minus donor-excluded-model weighted F1 on identical sites. Chromosome-level gains were averaged equally within each donor, then equally across retained donors. Models were not retrained.

**A. Overall chromosome-evaluation gains**

| Bin | Donors | Mean gain | 95% CI | Positive / N | p |
|---|---|---|---|---|---|
| 3 | 32 | +0.0381 | [+0.0296, +0.0466] | 29/32 | $2.56\times10^{-6}$ |
| 4 | 32 | +0.0280 | [+0.0206, +0.0359] | 28/32 | $1.93\times10^{-5}$ |

**B. Summary of exclusion checks**

| Bin | Omission | Mean-gain range | CIs above zero | Donors / check |
|---|---|---|---|---|
| 3 | One chromosome | +0.0377 to +0.0384 | 12/12 | 32 |
| 3 | One cell type | +0.0323 to +0.0437 | 6/6 | 24–28 |
| 4 | One chromosome | +0.0276 to +0.0285 | 12/12 | 32 |
| 4 | One cell type | +0.0261 to +0.0315 | 6/6 | 24–28 |

The mean-gain range gives the smallest and largest mean across the exclusion checks; it is not a confidence interval. All 32 donors remain in chromosome-omission checks. Cell-type omissions retain 24–28 donors, depending on the omitted group.

**Table A3 (continued). C. Leave-one-chromosome-out results**

| Bin | Omitted | Donors | Mean gain | 95% CI | p |
|---|---|---|---|---|---|
| 3 | Chromosome 6 | 32 | +0.0384 | [+0.0298, +0.0469] | $2.56 \times 10^{-6}$ |
| 3 | Chromosome 7 | 32 | +0.0381 | [+0.0295, +0.0466] | $2.56 \times 10^{-6}$ |
| 3 | Chromosome 8 | 32 | +0.0379 | [+0.0295, +0.0464] | $2.56 \times 10^{-6}$ |
| 3 | Chromosome 9 | 32 | +0.0383 | [+0.0297, +0.0469] | $2.56 \times 10^{-6}$ |
| 3 | Chromosome 10 | 32 | +0.0383 | [+0.0298, +0.0468] | $2.56 \times 10^{-6}$ |
| 3 | Chromosome 11 | 32 | +0.0382 | [+0.0296, +0.0467] | $2.56 \times 10^{-6}$ |
| 3 | Chromosome 12 | 32 | +0.0377 | [+0.0292, +0.0462] | $2.56 \times 10^{-6}$ |
| 3 | Chromosome 13 | 32 | +0.0381 | [+0.0296, +0.0467] | $2.56 \times 10^{-6}$ |
| 3 | Chromosome 14 | 32 | +0.0380 | [+0.0295, +0.0466] | $2.56 \times 10^{-6}$ |
| 3 | Chromosome 15 | 32 | +0.0381 | [+0.0296, +0.0467] | $2.56 \times 10^{-6}$ |
| 3 | Chromosome 16 | 32 | +0.0380 | [+0.0295, +0.0465] | $2.56 \times 10^{-6}$ |
| 3 | Chromosome 17 | 32 | +0.0382 | [+0.0296, +0.0467] | $2.56 \times 10^{-6}$ |
| 4 | Chromosome 6 | 32 | +0.0280 | [+0.0206, +0.0359] | $1.93 \times 10^{-5}$ |
| 4 | Chromosome 7 | 32 | +0.0280 | [+0.0206, +0.0359] | $1.93 \times 10^{-5}$ |
| 4 | Chromosome 8 | 32 | +0.0278 | [+0.0204, +0.0356] | $1.93 \times 10^{-5}$ |
| 4 | Chromosome 9 | 32 | +0.0285 | [+0.0210, +0.0363] | $1.93 \times 10^{-5}$ |
| 4 | Chromosome 10 | 32 | +0.0281 | [+0.0207, +0.0360] | $1.93 \times 10^{-5}$ |
| 4 | Chromosome 11 | 32 | +0.0281 | [+0.0207, +0.0359] | $1.93 \times 10^{-5}$ |
| 4 | Chromosome 12 | 32 | +0.0276 | [+0.0202, +0.0355] | $1.93 \times 10^{-5}$ |
| 4 | Chromosome 13 | 32 | +0.0279 | [+0.0204, +0.0357] | $1.93 \times 10^{-5}$ |
| 4 | Chromosome 14 | 32 | +0.0279 | [+0.0206, +0.0357] | $1.93 \times 10^{-5}$ |
| 4 | Chromosome 15 | 32 | +0.0281 | [+0.0208, +0.0360] | $1.93 \times 10^{-5}$ |
| 4 | Chromosome 16 | 32 | +0.0282 | [+0.0209, +0.0361] | $2.56 \times 10^{-6}$ |
| 4 | Chromosome 17 | 32 | +0.0282 | [+0.0208, +0.0361] | $2.56 \times 10^{-6}$ |

Each row omits the named chromosome from chromosomes 6–17. All 32 donors contribute to every chromosome-omission check.

**Table A3 (continued). D. Leave-one-cell-type-out results**

| Bin | Omitted | Donors | Mean gain | 95% CI | p |
|---|---|---|---|---|---|
| 3 | Bladder epithelial | 27 | +0.0359 | [+0.0265, +0.0454] | $4.92\times10^{-5}$ |
| 3 | Breast basal epithelial | 28 | +0.0364 | [+0.0272, +0.0458] | $2.74\times10^{-5}$ |
| 3 | Cortex neurons | 24 | +0.0323 | [+0.0231, +0.0414] | $2.77\times10^{-4}$ |
| 3 | Cardiomyocytes | 27 | +0.0391 | [+0.0292, +0.0488] | $4.92\times10^{-5}$ |
| 3 | Heart fibroblasts | 28 | +0.0437 | [+0.0364, +0.0514] | $7.45\times10^{-9}$ |
| 3 | Liver hepatocytes | 26 | +0.0407 | [+0.0305, +0.0507] | $8.80\times10^{-5}$ |
| 4 | Bladder epithelial | 27 | +0.0271 | [+0.0189, +0.0362] | $3.11\times10^{-4}$ |
| 4 | Breast basal epithelial | 28 | +0.0261 | [+0.0181, +0.0348] | $1.80\times10^{-4}$ |
| 4 | Cortex neurons | 24 | +0.0275 | [+0.0204, +0.0344] | $2.77\times10^{-4}$ |
| 4 | Cardiomyocytes | 27 | +0.0279 | [+0.0194, +0.0371] | $3.11\times10^{-4}$ |
| 4 | Heart fibroblasts | 28 | +0.0315 | [+0.0241, +0.0394] | $2.16\times10^{-7}$ |
| 4 | Liver hepatocytes | 26 | +0.0280 | [+0.0192, +0.0376] | $5.34\times10^{-4}$ |

Each row omits all donors of the named cell type. The cohort comprises bladder epithelial cells (5 donors), breast basal epithelial cells (4), cortex neurons (8), cardiomyocytes (5), heart fibroblasts (4), and liver hepatocytes (6).

For panels A, C and D, brackets give unadjusted 95% percentile donor-bootstrap confidence intervals (20,000 resamples). P-values are unadjusted exact two-sided donor sign tests for positive versus negative donor gains; ties are excluded. These tests assess directional consistency across donors, while the confidence intervals quantify uncertainty in the mean gain. Positive / N in panel A counts donors with gain > 0. Bin 4 pools the two highest source ranges before F1 calculation. These evaluations reuse the study donors and do not constitute independent-cohort validation.

**Figure A1. Class-specific and overall performance**

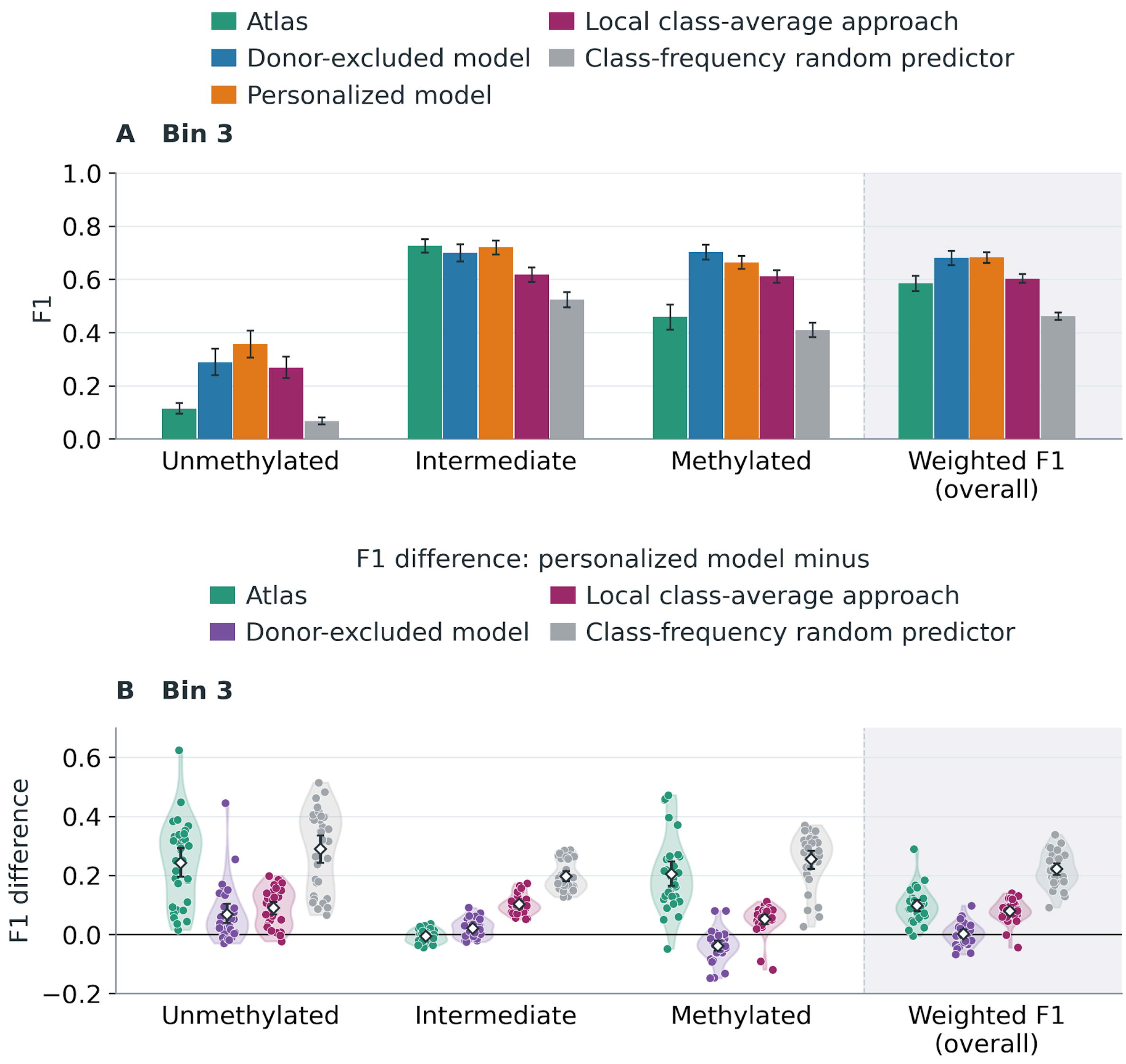


Figure A1. Class-specific and overall prediction performance across 32 donors. A–B: bin 3. A shows F1 averaged equally over donors for the atlas, donor-excluded model, personalized model, local class-average approach and class-frequency random predictor. B shows within-donor F1 differences between the personalized model and each reference. Dots represent donors and violins show their distributions. Diamonds mark mean differences; whiskers in both panels show 95% donor-bootstrap confidence intervals (20,000 resamples).

**Figure A1 (continued)**

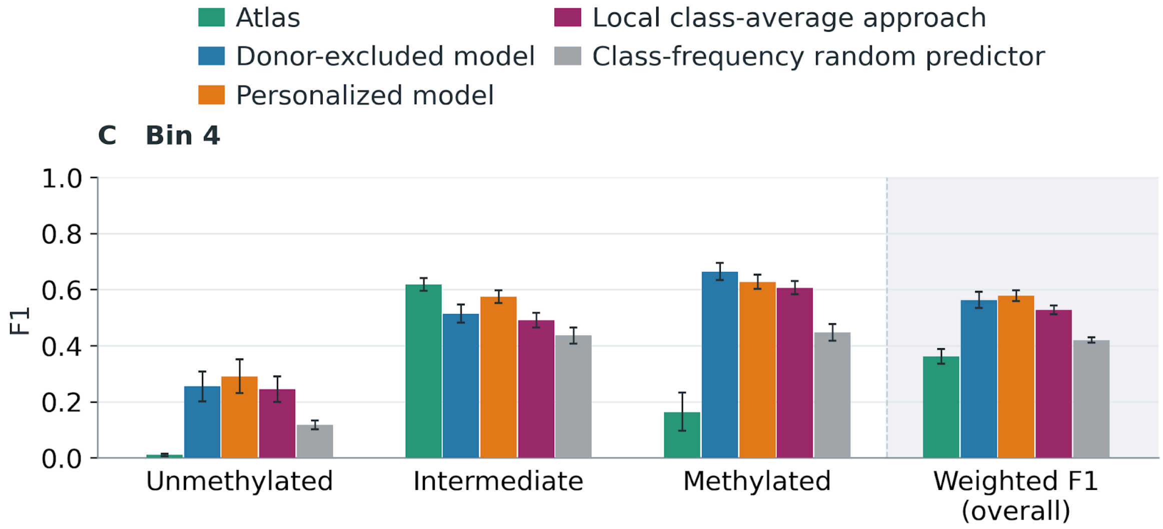


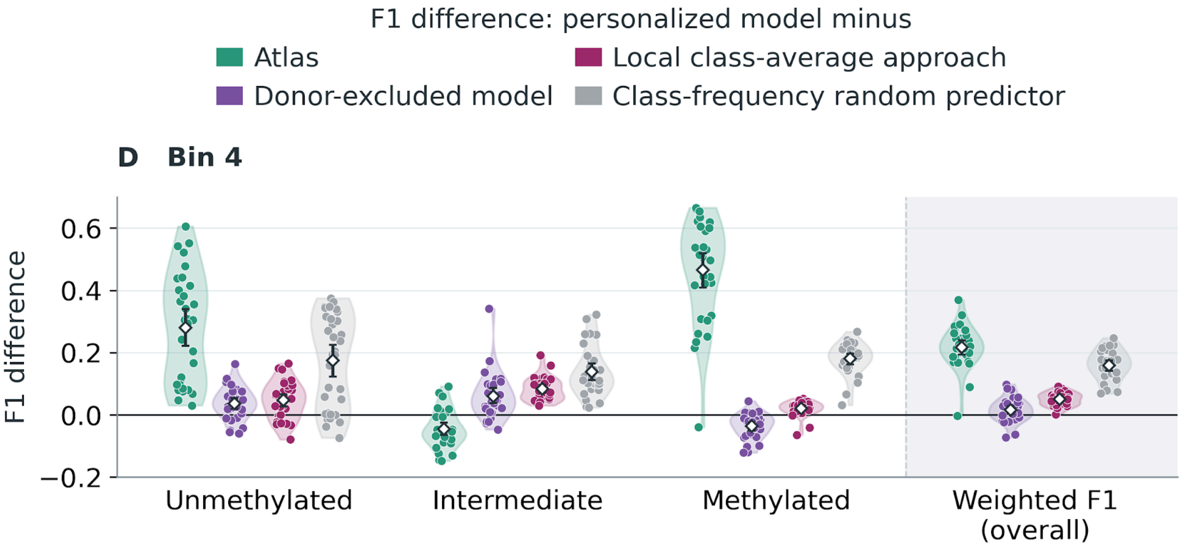


Figure A1 (continued). C–D: bin 4; plotting conventions are as in A–B. Purple denotes personalization gain relative to the donor-excluded model; magenta denotes the difference relative to the local class-average approach; green and gray denote differences relative to the atlas and random predictor. Overall weighted F1 weights classes by their true frequencies within each donor before averaging over donors. The random predictor is calculated from expected confusion counts using the evaluated donor's class frequencies. The atlas uses the target-label thresholds, including methylated ≥0.8.
